# Improved long read correction for de *novo* assembly using an FM-index

**DOI:** 10.1101/067272

**Authors:** James M. Holt, Jeremy R. Wang, Corbin D. Jones, Leonard McMillan

## Abstract

Long read sequencing is changing the landscape of genomic research, especially *de novo* assembly. Despite the high error rate inherent to long read technologies, increased read lengths dramatically improve the continuity and accuracy of genome assemblies. However, the cost and throughput of these technologies limits their application to complex genomes. One solution is to decrease the cost and time to assemble novel genomes by leveraging “hybrid” assemblies that use long reads for scaffolding and short reads for accuracy. To this end, we describe a novel application of a multi-string Burrows-Wheeler Transform with auxiliary FM-index to correct errors in long read sequences using a set of complementary short reads. We show that our method efficiently produces significantly higher quality corrected sequence than existing hybrid error-correction methods. We demonstrate the effectiveness of our method compared to state-of-the-art hybrid and long-read only *de novo* assembly methods.

## 2 Introduction

*De novo* genome assembly has benefitted dramatically from the introduction of so-called “long” read sequencing technologies. These technologies, such as SMRT sequencing by Pacific Biosciences and nanopore sequencing platforms by Oxford Nanopore Technologies, produce reads typically 10s of kilobases instead of hundreds of bases. These reads can span repetitive or low-complexity regions of the genome previous unresolvable using only “short”-read next-generation sequencing. Unfortunately, the relatively high error rate of these long-read technologies introduces new informatics and analysis challenges. Effective and efficient methods are necessary to correct these errors in order to realize the potential of these long reads for whole genome assembly [10, 2, 16, 22].

Long read correction algorithms can be broadly classified as either self-correction or hybrid correction algorithms. Self-correction algorithms correct long reads using only other long read sequences. Self-correcting algorithms, including Sprai [20], HGAP [10], and PBcR [16] align the long reads to each other and generate a consensus sequence. In order to generate an accurate consensus, these methods requires relatively high coverage of long read sequence to overcome the high error rate. Unfortunately, the relatively high cost per *accurate* nucleotide for long-read sequencing technologies often makes deep sequencing using only long reads expensive.

In contrast, hybrid correction algorithms use short-read sequencing of the same sample to complement and correct the long reads. Short-read sequencing has fewer sequencing errors, costs less per base sequenced, and thus the cost per *accurate* nucleotide is much lower. State-of-the-art hybrid correction algorithms include LoRDEC [22], Cerulean [26], ECTools [14], DBG2OLC [9], and hybridSPAdes [1]. These hybrid methods are often able to construct more accurate and contiguous assemblies than exclusively long-read assembly methods at substantially less cost.

For either class of method to be useful for large, complex genomes that are biomedically or economically important, the key challenge is performing as accurate as an assembly as possible in as short of time and with the least computational resources. Current methods often take hundreds to thousands of hours on high performance computing clusters with access to many nodes with large memory configurations[4, 22]. Given that finding the appropriate parameters for an assembly is often an iterative process, these high computational costs are a barrier.

We introduce a new hybrid method for correcting errors in long-read sequences called FM-index Long Read Corrector (FMLRC) that corrects long reads using a multi-string Burrows-Wheeler Transform (BWT) variant that has been adapted to string collections [3] and FM-index of a short-read sequencing dataset as an *implicit* de Bruijn graph [6] (see Figure 4). FMLRC can use a variety of input reads and does not require preassembly of the short reads. In brief, we construct a multi-string BWT from a set of short high-accuracy reads such as Illumina sequence. This multi-string BWT allows for both the compression of data and combining of multiple data sets [15].

This BWT is used as an implicit de Bruijn graph to implement a seed-and-extend or seed-and-bridge strategy analogous to that used in LoRDEC [22]. LoRDEC first generates a de Bruijn graph composed of *k*-mers from the short reads [22]. Then, the graph is pruned such that any low-frequency *k*-mers (specified by a user-defined threshold) are removed from the graph. Long reads are then compared against this graph and broken into regions that are labeled as either *solid* or *weak*. All *k*-mers within solid regions are contained in the pruned short-read de Bruijn graph. All *k*-mers within weak regions are not in the de Bruijn graph. In general, the assumption of LoRDEC is that weak regions are caused by errors in sequencing and should be replaced with the closest series of solid, overlapping *k*-mers from the de Bruijn graph. When weak *inner* regions are identified in a long read, the flanking solid *k*-mers are used as endpoints for finding a *bridge* (or path) in the de Bruijn to connect the two solid regions. In the event of multiple supported bridges, the one with the closest edit distance to the original sequence is chosen. The *head* and *tail* of the graph are symmetrical special cases where there is only one flanking solid region. In either case, LoRDEC searches for the best extension of the single solid region that most closely matches the weak head or tail sequence.

While LoRDEC has been shown to correct a large fraction of errors in long read sequences [22], the user must select a fixed, short *k*-mer size and a fixed threshold for pruning. The use of an explicit de Bruijn graph fundamentally limits the ability of LoRDEC to resolve repetitive or low-complexity elements longer than *k*. In the de Bruijn graph, low complexity sequences tend to look like “hairballs” of interconnected nodes where there are too many possible paths to explore. When LoRDEC enters a low complexity region, it will usually fail to find a path because it reaches a self-imposed limit on the graph exploration. Additionally, parameters are often chosen heuristically and changing them requires re-computing the entire de Bruijn graph prior to re-running the correction.

In contrast, FMLRC finds *k*-mer “seeds” in the long read sequence with high support in the BWT, then searches for a high-weight path between seeds that most closely matches the intervening long read sequence. Multiple correction passes using increasing anchor sizes, k, allows us to resolve small-scale errors while avoiding inaccurate de Bruijn graph traversals caused by repetition of shorter *k*-mers. Since the FM-index can search for substrings of any length, our method is not constrained to a single fixed *k*-mer size, so it represents all possible de Bruijn graphs for the short-read sequencing data. Additionally, the BWT is a lossless encoding of the short reads, allowing any pruning threshold to be dynamically adjusted without needing to reconstruct an entire de Bruijn graph. Our method is unique in that it applies both a short *k*-mer and long *K*-mer de Bruijn graph to the correction process, allowing for the correction algorithm to correct through low complexity regions up to the size of the long *K*-mer. An overview of our approach is shown in Figure 1.

**Figure 1:**
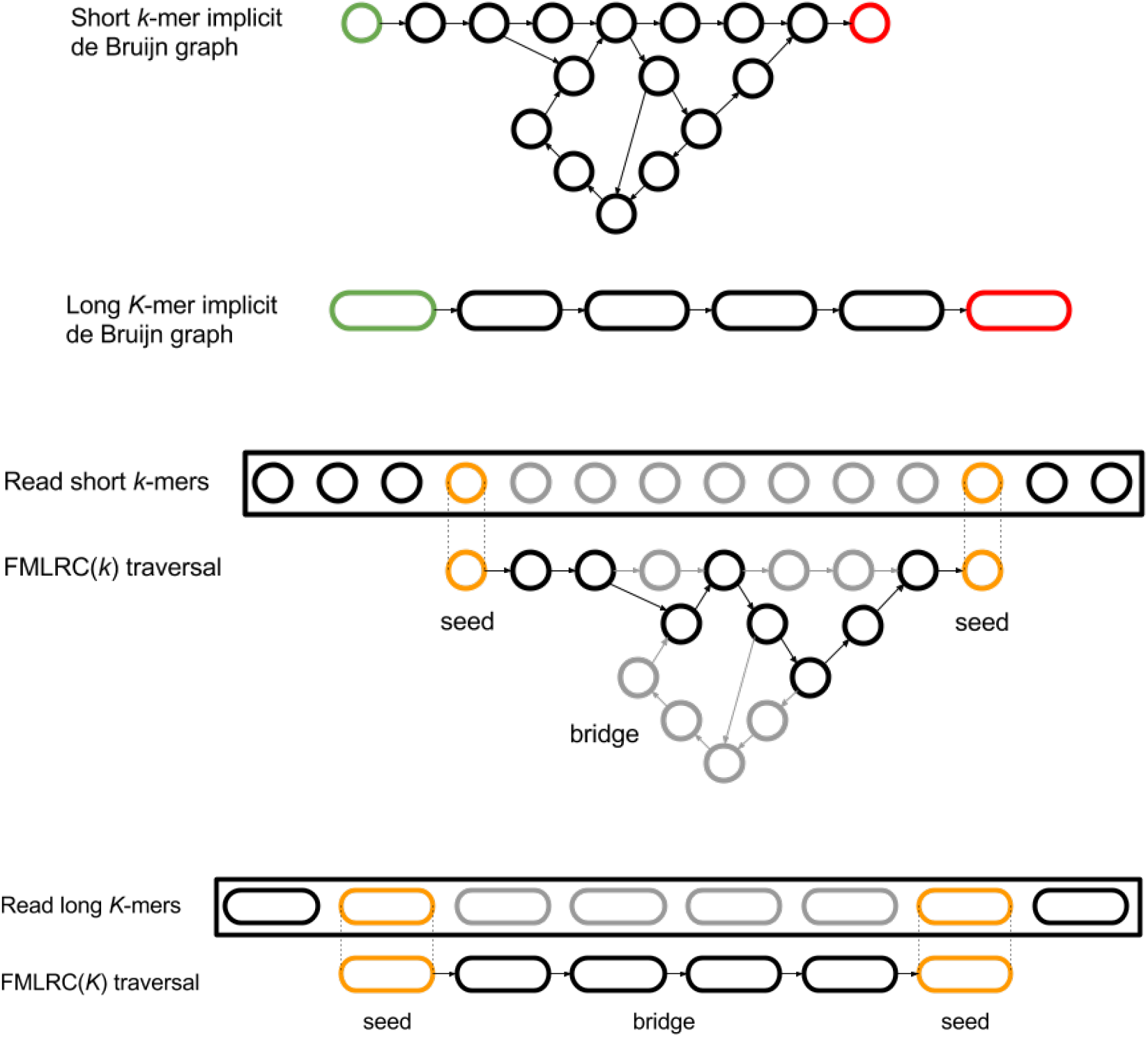
Illustration of the seed-and-bridge correction strategy using short and long *k*-mers. *Implicit* de Bruijn graphs with arbitrary *k* can be inferred from a multi-string BWT. The use of a short, fixed *k* often does not resolve “hairball” and other structures in the graph caused by low-complexity and repetitive genomic elements. Longer *K*-mers may dramatically simplify the bridging step if sufficiently long seeds can be found. Illustrative seed-and-bridge paths are shown for short *k*-mer and long *K*-mer graphs. Seed *k*-mers are shown in orange, and the correct path in black. The two-pass (*k, K*) seed-and-bridge correction implemented in FMLRC allows the correction of short, nonrepetitive segments in the first pass, then seeding larger *K*-mers and bridging to resolve more complex sequences.

As a hybrid method for correcting errors in long-read sequences, the key advantage of FMLRC is its use of a BWT with FM-index allows iterative correction of errors in sequences of arbitrary length by constructing paths through an implicit de Bruijn graph. The flexibility of *k*-mer nodes in the graph allows us to resolve low-complexity and repetitive elements more efficiently and with greater accuracy than existing hybrid error-correction methods. We illustrate this by comparing the assembly of long reads using modern overlap-layout assemblers to FMLRC.

## 3 Results

### 3.1 Evaluating computational efficiency and accuracy

A key difference in FMLRC is that it performs two passes of correction: a relatively short *k*-mer pass followed by a longer *K*-mer pass. In each pass, the “pruning” threshold that determines if a *k*-mer is solid or weak is dynamically adjusted based on the *k*-mer length and the surrounding *k*-mer frequencies in the read. The first pass performs the majority of error correction and is conceptually similar to other correction methods based on de Bruijn graphs. The second pass using the longer *k*-mer allows FMLRC to assemble through low-complexity or repeat regions in the reads that other algorithms are not capable of easily assembling through. To demonstrate the effectiveness of this approach in FMLRC, we evaluated the accuracy of our method using complementary long- and short-read datasets for five species: *E. coli K12, P. falciparum 3d7, S. cerevisiae W303*, and *A. thaliana* (see Section 5.9). We performed a detailed comparison between our method and LoRDEC to show the relative correction accuracy and computational performance. We then assessed the effectiveness of our corrected reads for *de novo* assembly using a non-correcting assembler, miniasm [18], and compared these data to several other state-of-the-art hybrid and long-read-only *de novo* assembly methods.

### 3.2 Accuracy of corrected reads

We ran FMLRC to correct the long reads in these datasets and compared the corrected read accuracy to that produced by LoRDEC. We assessed LoRDEC alone (LoRDEC(*k*)), LoRDEC followed by FMLRC’s second pass (LoRDEC(*K*)+FMLRC(*K*)), and the full two-pass FMLRC (FMLRC(*k, K*)). In all cases, either LoRDEC(*k*)+FMLRC(*K*) or FMLRC(*k, K*) had the highest accuracy (Figures 2a and 2b). The second (long *K*) pass of FMLRC strictly improved the results of LoRDEC(*k*) by increasing the number of matching bases and percent matching. Thus, this shows that a two pass approach in general provides better correction, regardless of which algorithm was used for the short *k*-mer pass. Second, it suggests that for many genomes, but not all, FMLRC *k*-mer pass provides somewhat better correction.

**Figure 2:**
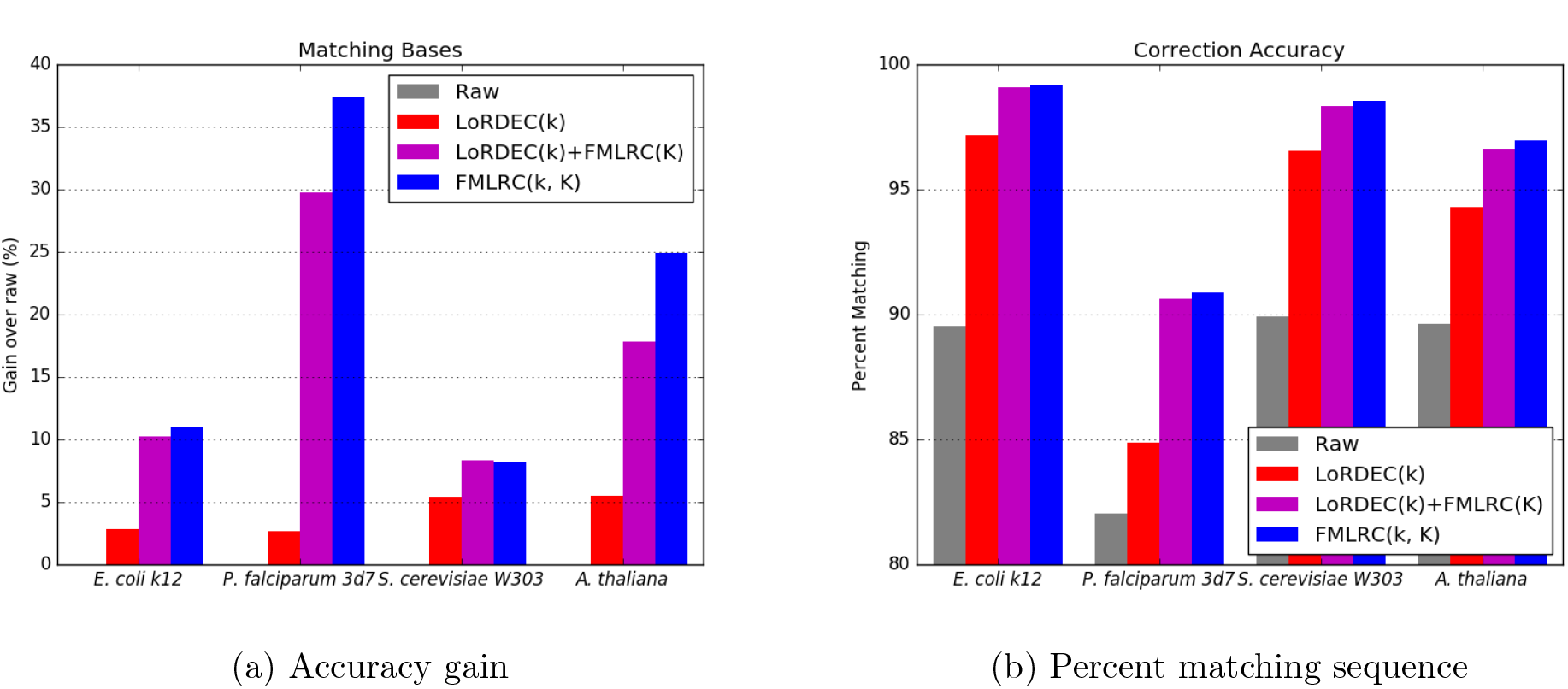
After aligning the corrected reads, we counted the number of bases that exactly match the reference mapping. We then normalized each value by the number of bases that matched the reference in the raw uncorrected reads and plotted the value as a percentage gained (a). Note that the raw reads always have a gain of 0%. In all test cases, the reads corrected by FMLRC have more matching bases than than those from only LoRDEC, indicating that FMLRC is corrected more bases to match the reference. We also calculated the fraction matching of each alignment by taking the total number of matching bases and dividing by the total number of aligned bases. We plotted the results as a percentage of aligned bases that match (b). In all test cases, FMLRC has a higher fraction mapping than LoRDEC.

### 3.3 Performance

As the size of long read datasets and genomes undergoing *de novo* assembly increases, the performance of hybrid long read correction and assembly methods becomes increasingly important. For genomes of more complex eukaryotes and mammals, the computational resources required for effective *de novo* assembly are staggering and difficult to coordinate. This is driven largely by the pairwise overlap step required by all modern long read assemblers. The time required to overlap these long reads with one another increases quadratically relative to the number of reads. While novel methods such as MHAP [4] and minimap [18] aim to improve this, in practice, the computational time and memory required are often prohibitively expensive. Hybrid error correction methods that correct long reads without pairwise overlapping dramatically simplify the subsequent overlap and layout of long reads for assembly by reducing the variance that must be accounted for in the overlapping step. In particular, long reads having undergone error correction are likely to share much longer identical stretches that can be used to efficiently find confidently overlapping reads. Fundamentally, the longer and more accurate these corrections are, the more quickly and accurately the long reads can be assembled. The majority of long read error correction methods act similar to scaffolders in that they require the assembly of complementary short read data first, then alignment between long reads and short-read unitigs or contigs. These approaches, while reasonably effective, suffer from two classes of problems. First, they incur the same type of disadvantages a short-read only assemblies in that low-complexity and repetitive elements larger than the size of the short reads cannot be reliably resolved. When short reads are preassembled, this bias can “correct” long read with incorrect sequence, confounding assembly. Second, the short read assembly and pairwise alignment/overlap of long reads with short-read contigs incur performance penalties approaching those of full pairwise long-read overlapping.

As previosuly shown [22], use of an explicit (LoRDEC) or implicit (FMLRC) de Bruijn graph to implement a seed-and-extend or seed-and-bridge strategy is expected to be computationally efficient compared to other methods. To confirm this expectation, we assessed the CPU time required for LoRDEC(*k*), the short read BWT construction, FMLRC(*K*), and FMLRC(*k, K*) for four different genomes of varying complexity. The BWT construction was performed using a combination of *ropebwt2* [17] and the *msbwt* package^1^. Note that the BWT construction is a pre-process that is computed *once* per short-read dataset prior to to any FMLRC execution. FMLRC(*K*) is a single pass using only a long *K*-mer that is run on the result of LoRDEC(*k*). FMLRC(*k, K*) is the implementation of the two-pass algorithm we describe.

For the *E. coli* and *S. cerevisiae* datasets, FMLRC(*k, K*) requires less CPU time than LoRDEC(*k*) (Figure 3). For the *P. falciparum* and *A. thaliana* datasets, the two pass FMLRC(*k, K*) requires more CPU time, but also has the largest relative gains in correction performance over LoRDEC(*k*) in these two tests.

**Figure 3:**
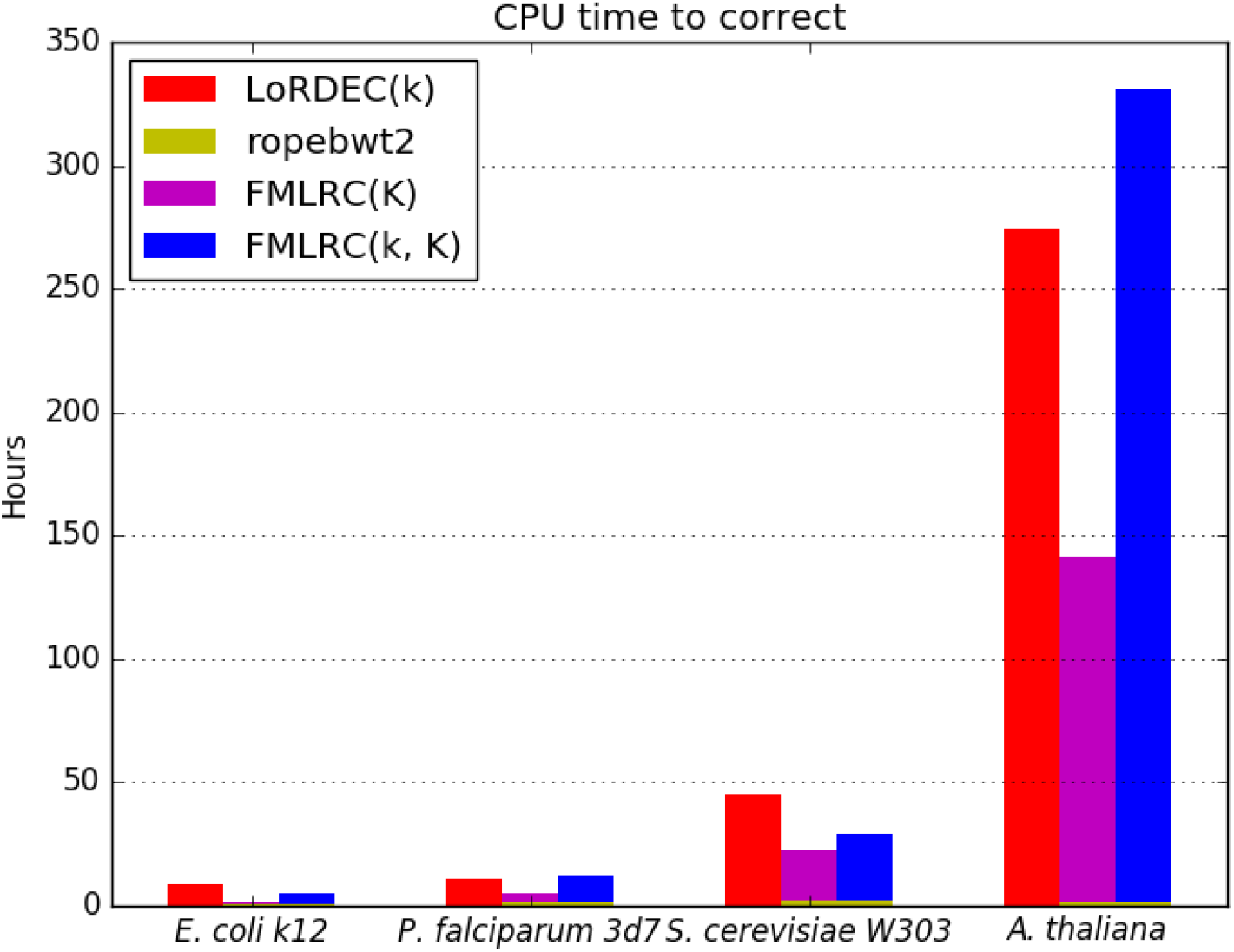
CPU time. This figure shows the CPU time required by LoRDEC and FMLRC. Ropebwt2 is a BWT construction algorithm that is run once prior to any executions of FMLRC, so it is shown stacked with both versions of FMLRC. FMLRC(K) was run on the results of LoRDEC(k).

### 3.4 *De novo* Assembly

The ultimate goal of any long read correction algorithm is to provide better data for genomics and metagenomic analysis. We assessed the ability of our method to successfully complete assembly of simple and complex genomes and to compare its performance to other leading long-read error correction and *de novo* assembly methods shown in Table 1. We performed assemblies of three of the genomes (*E. coli K12, S. cerevisiae W303*, and *A. thaliana*) using all of these methods.

**Table 1:** Long-read and hybrid correction and assembly methods. All of the compared methods are shown along with their mode of error correction and assembly, each either long-read only or “hybrid” using complementary short-read data. “Preassembly” indicates whether a hybrid method requires the short read data to be preassembled using a different method.

| Method | Correction | Assembly | Preassembly | Citation |
| --- | --- | --- | --- | --- |
| miniasm |  | Long-read |  | [18] |
| canu | Long-read | Long-read |  | [4] |
| Sprai | Long-read |  |  | [20] |
| hybridSPAdes |  | Hybrid |  | [1] |
| DBG2OLC |  | Hybrid | X | [9] |
| Cerulean |  | Hybrid | X | [26] |
| ECTools | Hybrid |  | X | [14] |
| LoRDEC | Hybrid |  |  | [22] |
| FMLRC | Hybrid |  |  | Our method |

Our method, along with LoRDEC, ECTools, and Sprai, perform only read correction. We used miniasm to assemble the corrected reads from these methods. The straightforward approach to identity-based overlapping and graph layout used by miniasm allows us to assess the effect of read correction on *de novo* assembly.

Canu is a modern fork of the Celera Assembler and consists of the basic PBcR correction method using the MHAP overlapper followed by assembly with HGAP. So we assess only the canu pipeline as a whole.

Several of the methods took prohibitively long or failed to assemble the *A. thaliana* genome. We analyzed completed assemblies using Quast [13] in Table 2. Percent error indicates the total of mismatched bases, insertions/deletions, and no-calls (Ns). As described by [13], NGA50 is analogous to N50, but breaks misassembled contigs and is taken as a percentage of the real genome size instead of the assembly size. As shown, FMLRC has comparable performance to other methods for *E. coli K12*. It also outperforms all methods tested in terms of N50 and all but Cerulean in terms of NGA50 for *S. cerevisiae W303*. Although the error rate is lower for canu and hybridSPAdes, these typically rely on exceptionally high coverage of long reads and short reads, respectively, and degrade in performance quickly as coverage drops. These test datasets contain high (>100x) coverage of both long and short reads. Furthermore, post-assembly polishing steps such as Quiver [10] and Nanopolish [19] are typically effective in reducing the assembly error from less than 1% to less than 0.01%.

**Table 2:**
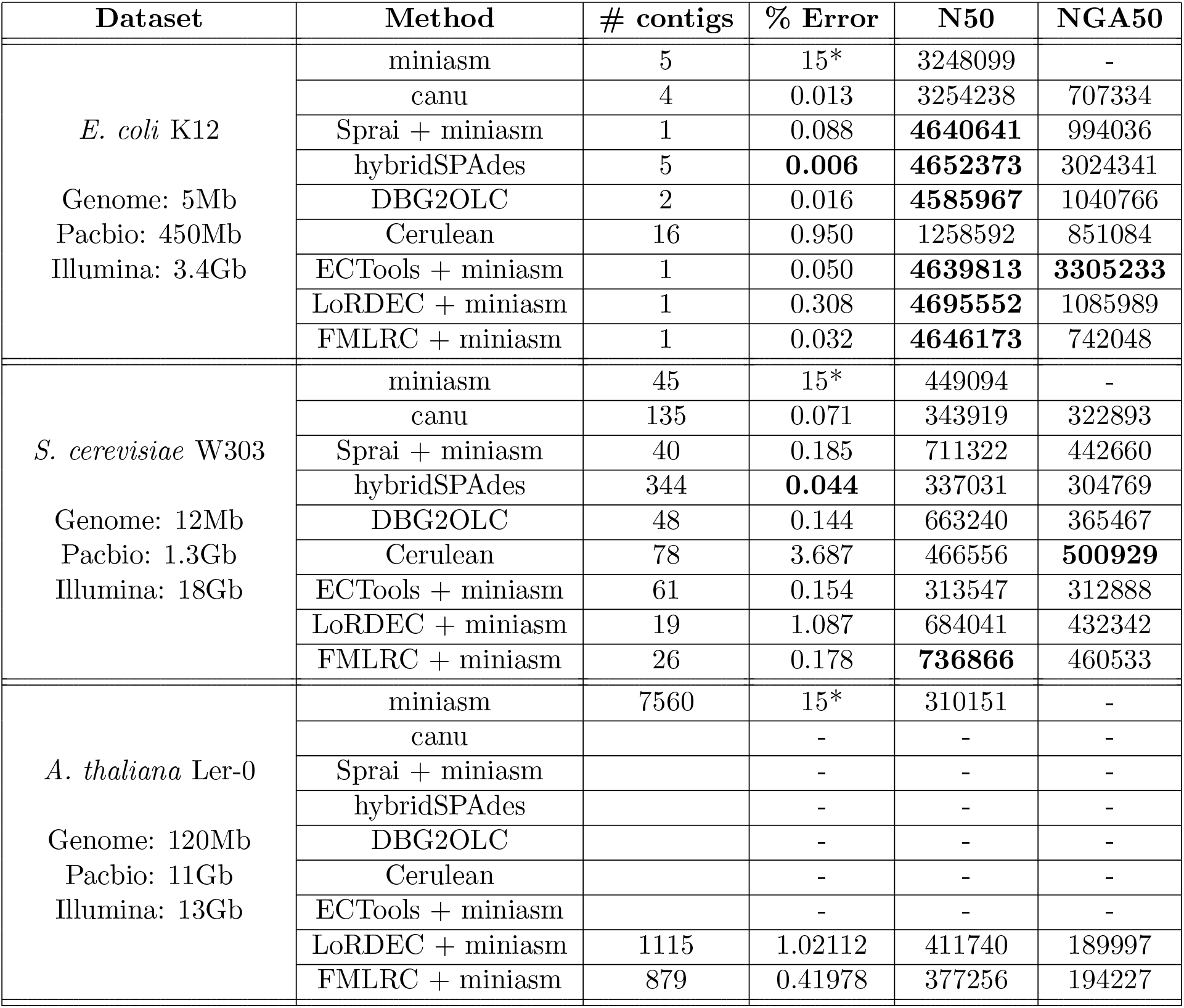
Long-read and hybrid correction assembly statistics. For *A. thaliana*, the methods marked with dashes did not complete in under a week with 16 processes. *miniasm does not perform either read correction or consensus calling, so the resulting assembly has the same error profile of the input reads.

Our conclusion is that read correction by FMLRC before overlapping increases the sensitivity and/or specificity of long-read overlapping and layout leads to a more contiguous assembly.

## 4 Discussion

Flexible “modular” approaches to *de novo* long read sequence assembly are becoming more popular with the introduction of efficient overlap and layout methods such as DALIGNER [21], MHAP [4], minimap [18], and miniasm [18]. Existing error correction methods including DBG2OLC [9], Cerulean [26], hybridSPAdes [1], and ECTools [14] require preassembly of short read sequence and perform a variant of scaffolding using long read sequences. While this approach benefits from the high accuracy of short read sequence, it retains the biases inherent in assembly of short read sequences. In particular, it is often difficult or impossible to properly assemble low-complexity or repetitive sequences using only short reads [25].

To overcome these limitations, we have introduced FMLRC, a long read correction method that uses a multi-string BWT and FM-index as an implicit de Bruijn graph. The method uses two passes to perform the correction: one with a relatively short *k*-mer and one with a longer *K*-mer. In each pass, the method searches for weak, unsupported regions in the reads and uses the implicit de Bruijn graph to identify alternate solid paths to correct the reads. The BWT with FM-index serves as an efficient lossless compression of read datasets and allows more flexible and thorough navigation of the short read sequence. Without modification, the MSBWT implicitly represents de Bruijn graphs with arbitrary *k*, supporting the efficient multi-pass correction method in FMLRC.

We compared the results of FMLRC to the LoRDEC algorithm for performing hybrid error correction. We showed that our method reliably produces higher accuracy corrections than LoRDEC and is computationally efficient. We further showed that using FMLRC as a preassembly error correction step in conjunction with existing overlap-layout assembly methods produces highly contiguous assemblies with competitive accuracy relative to existing hybrid and non-hybrid assembly methods.

Future work will include a specific cost-benefit analysis of the quantity of long- and short-read data required to effectively assemble genomes based on their size and repetitive structure. While previous work has been done in this area, FMLRC, as a more efficient method for hybrid correction of long reads, is expected to allow more effective *de novo* assembly with less long read data than previously possible. In the long term, better integration of FMLRC error correction along with other tools for overlapping, layout, and consensus of long read sequencing data will help realize the goal of a fully modular and efficient *de novo* assembly process.

## 5 Methods

The FM-index Long Read Corrector (FMLRC) is a hybrid correction method that uses the BWT [7] and FM-index [11] of a short-read sequencing dataset to correct a long-read sequencing dataset. These data structures have been previously used for short-read self-correction in FMRC [12], but the greedy method of correction used by FMRC is ill-suited for long-read correction.

The major difference between FMLRC and other hybrid correction algorithms is the use of a BWT and FM-index of the short-read sequencing dataset as a de Bruijn graph [6]. The BWT and FM-index have several advantages over a typical de Bruijn graph. First, the BWT is a lossless encoding of the original reads, meaning that no reads (or *k*-mers) are “pruned” as they commonly are in a de Bruijn graph. Secondly, the FM-index enables arbitrary *k*-mer frequency lookups in O(*k*) steps. Thirdly, the FM-index is not fixed a single value of k, allowing methods to dynamically choose a *k*-mer value at run-time. The combination of these three properties means the BWT and FM-index implicitly represent *all* de Bruijn graphs for the short-read sequencing dataset.

Given that the BWT and FM-index implicitly represent all de Bruijn graphs, FMLRC uses the graph to perform long-read correction. Each long read is individually broken into *solid* and *weak* regions. *Solid* regions are composed of *k*-mers supported by the implicit de Bruijn graph at a threshold and *weak* regions are not supported at that threshold. Given two solid regions with a single weak region in between, the algorithm searches for solid *bridges* in the de Bruijn graph that connect the solid regions and span the weak regions. If no bridges are found, no change is made to the long read. If one or more bridges are found, they are aligned to the original weak region, and the one with the smallest edit distance is chosen to replace it. Weak regions at the beginning or end of the long read are treated similarly, but with a seed-and-extend assembly as opposed to a seed-and-bridge method.

The key difference in FMLRC is that it performs two passes of correction: a relatively short *k*-mer pass followed by a longer *K*-mer pass. In each pass, the “pruning” threshold that determines if a *k*-mer is solid or weak is dynamically adjusted based on the *k*-mer length and the surrounding *k*-mer frequencies in the read. The first pass performs the majority of error correction and is conceptually similar to other correction methods based on de Bruijn graphs. The second pass using the longer *K*-mer allows FMLRC to assemble through low-complexity or repeat regions in the reads that other algorithms are not capable of easily assembling through.

### 5.1 Correction of Long Read Sequences using BWTs

Our method assumes that a BWT and FM-index of the short reads has been constructed. The BWT construction described here was performed using a combination of *ropebwt2* [17] and the *msbwt* package^2^. As a BWT and FM-index is not restricted to any particular *k*-mer length, it acts as an implicit un-pruned de Bruijn graph for both *k* and *K* such that the frequency of any *k*-mer or *K*-mer in the dataset can be looked up in respectively *O(k)* or *O(K)* steps.

For our method, the two passes are programmatically identical with the value of *k* or *K* passed in as a parameter. For brevity, we describe the FMLRC correction using parameter *k* noting that replacing *k* with *K* describes the second pass of our method. Additionally, all thresholds and parameters are described as functions of *k* and the implicit *k*-mer de Bruijn graph.

### 5.2 The BWT and Implicit De Bruijn Graph

The BWT and FM-index implicitly represent all de Bruijn graphs for the short-read dataset. When a node in any of the graphs needs to be accessed, FMLRC uses the FM-index to perform a *k*-mer search in *O(k)* steps. This informs FMLRC of the *k*-mer frequency in the dataset. FMLRC then compares the *k*-mer frequency to a pruning threshold and classifies it as *weak* if it is below the threshold and *strong* if it is above the threshold. In this regard, the BWT and FM-index are advantageous data structures because values such as *k*-mer size and the pruning threshold distinguishing solid and weak can be dynamically adjusted before or during the correction process without needing to explicitly recompute the full de Bruijn graph.

FMLRC includes two FM-index implementations. The default FM-index implementation uses bit arrays and rank operations to enable fast *k*-mer lookups at the cost of a larger memory footprint.

The second FM-index implementation is a traditional sampled FM-index that allows users to set the sampling rate, leading to longer computations with a smaller memory footprint. The two FM-index implementations produce identical corrected read results, and we present only the results from the default implementation in our performance results.

### 5.3 Identification of Weak Regions

Given a long read and a BWT of the short reads, the first step is to identify weak regions of the long read. To do this, our method uses a hybrid threshold that is a function of *k*-mer length and *k*-mer frequency in surrounding solid *k*-mers. First, the method retrieves *k*-mer counts for every *k*-mer in the long read. For most long reads, the majority of *k*-mer counts are at or near zero due to the high error rate from the sequencing technology. Second, every *k*-mer with a count that is less than a user-defined, absolute minimum threshold, *T*, is removed from the list of counts. Next, the median, m, of the remaining counts (that are all ≥ *T*) is calculated. A second user-defined value, *F*, is the fraction of the median m that is required for a path to be consider solid. Finally, the solid threshold *t* for a particular read is calculated as *t* = max(*T, F * m*). In other words, the method calculates a dynamic threshold based on *k*-mer counts from that particular read. However, if that dynamic threshold is below an absolute, user-defined minimum, the absolute minimum is used as the threshold instead. For low-coverage short-read datasets, it is often the case that *t* = *T* because *F * m < T*. For high-coverage short-read datasets, this dynamic threshold alleviates the need to select a fixed threshold beforehand, and it instead uses counts from the implicit de Bruijn graph to derive an expected count for *k*-mers in the read. Given the read-specific threshold *t*, weak regions are identified as contiguous weak *k*-mers (the frequency of each *k*-mer is < t). Similarly, solid regions are contiguous solid *k*-mers (the frequency of each *k*-mer is ≥ *t*).

### 5.4 Correction of Weak Regions

Weak regions can be flanked by zero, one, or two solid regions. If a weak region has no flanking solid regions, the entire read is one large weak region with no solid *k*-mers to initialize a traversal of the de Bruijn graph. As a result, these reads are simply written to the output with no changes because there are no solid seeds.

If a weak region has one flanking solid region, then it is either the head or tail weak region for the read. In either case, the solid *k*-mer closest to the weak region is used as the initializer *k*-mer for traversing the de Bruijn graph using a seed-and-extend approach. The de Bruijn graph is explored from this start *k*-mer using a depth-first traversal and an expected path length based on the distance to the end of the read. Additionally, our method enforces a limit on the amount of branching that is allowed to reduce computation time. This branch limit is usually only reached if the algorithm is traversing a complex section of the de Bruijn graph where many interconnected nodes lead to an exponential number of graph traversals. If no paths are found or the branch limit is reached, then no change is made to the read. If one or more paths are found, then they are each aligned to the original weak head/tail region and the one with the smallest edit distance replaces the original.

Finally, the most common case is a weak region flanked by two solid regions. In this case, we use a seed-and-bridge approach by selecting an initial starting *k*-mer is selected from the first solid region and a target *k*-mer is selected from the second. The de Bruijn graph is explored in a depth-first traversal from the starting *k*-mer. Thus, the algorithm attempts to find a path in the de Bruijn graph that bridges the two solid regions and spans the weak region. Each bridge is extended so long as it is shorter than an expected length that is estimated from the distance between the two solid *k*-mers in the long read. Whenever a bridge is found that ends with the target *k*-mer, it is added to a list of discovered bridges. Similar to the head and tail traversal, the bridge traversals also have a branch limit to reduce the amount of computation. If no bridges are found or the branch limit is reached, then no change is made to the read. If one or more bridges are found, then they are each aligned to the original weak region and the one with the smallest edit distance is used to replace the original.

### 5.5 Differences in the Short and Long Passes

FMLRC is a two pass algorithm using first a short *k*-mer followed by a second longer *K*-mer. In general, the short *k*-mer pass does the majority of the correction for the method. While the *k*-mers are described as “short”, they are still long enough to uniquely identify most sections of the genome. If a section can be uniquely identified using short *k*-mers, then correction is often a relatively easy process and a longer *K*-mer is not required. This is why many other *k*-mer based methods are able to correct the majority of errors in long reads.

The short *k*-mer pass is conceptually similar to other previously described methods that use a de Bruijn graph for correction such as LoRDEC [22]. However, these previous methods are restricted to a fixed *k*-mer size, fixed pruning threshold, and a fixed branching limit; all of which are typically user-defined parameters that are commonly chosen heuristically. Despite these limitations, it is worth noting that these other correction methods can be used in place of the first *k*-mer pass of FMLRC. Thus, the full pipeline could be described as a short *k*-mer correction method followed by our long *K*-mer correction pass. In our results, we test FMLRC as both a two-pass method and as a one-pass long *K*-mer method following a different short *k*-mer correction method (LoRDEC [22]).

To the best of our knowledge, the long *K*-mer pass is unique to FMLRC. To provide some intuition behind why it improves the results, we focus on the differences in de Bruijn graphs representing the same data but with two different *k*-mer lengths. In general, two distinct paths will be merged in a *k*-mer de Bruijn graph if they share a pattern that is at least *k* long. This is because the nodes along that shared region will be identical. At the ends of the shared region, there will be two paths emerging representing the differences at the edge of the shared regions. For example, Figure 4a shows an example de Bruijn graph using short 3-mers to represent two strings.

**Figure 4:**
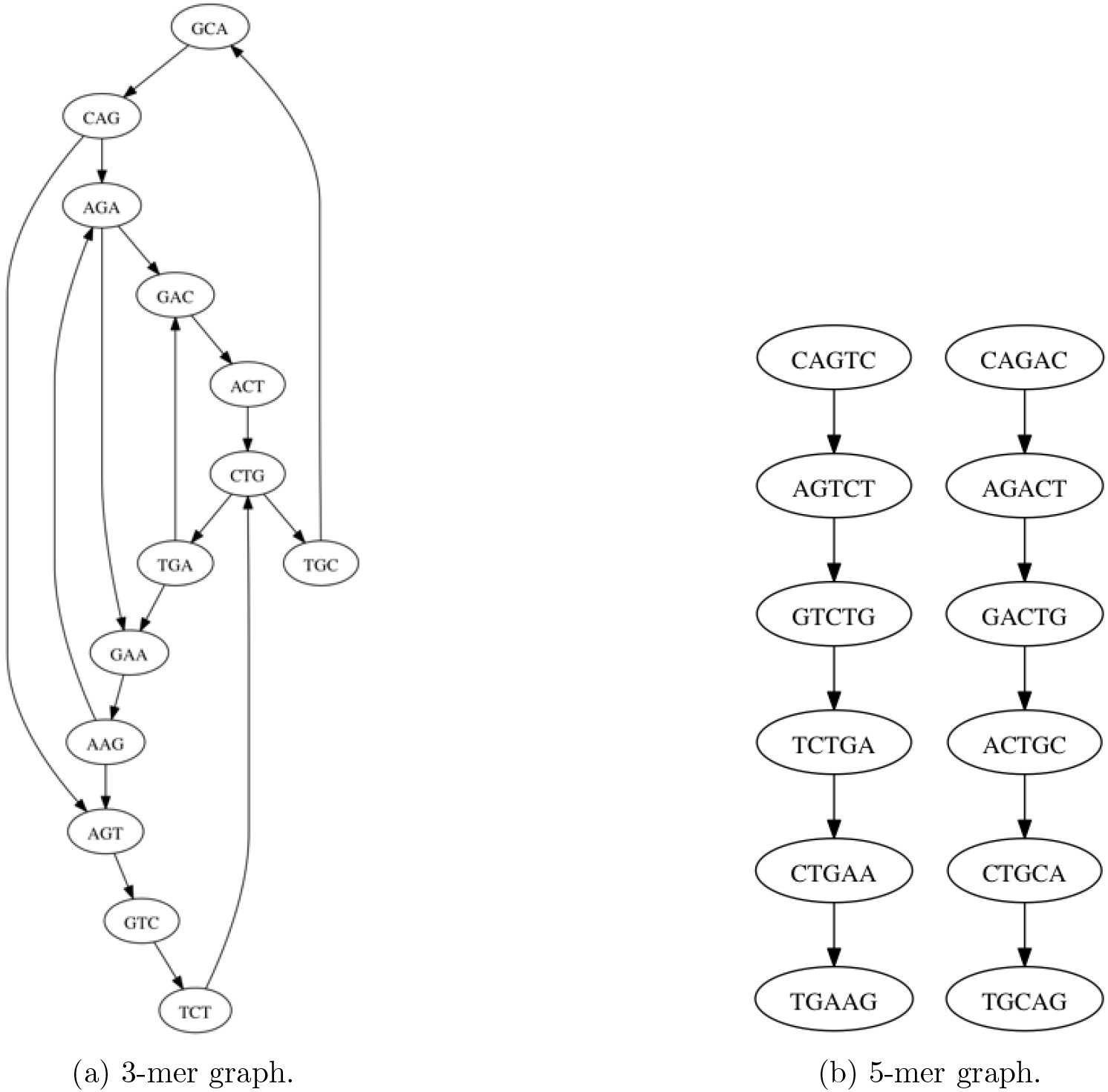
Two de Bruijn graphs for strings “CAGACTGCAG” and “CAGTCTGAAG” are shown. A 3-mer graph is shown in (a), where every 3-mer from the two strings is a node. An edge indicates that the 2-mer suffix of the source 3-mer is the same as the 2-mer prefix of the destination 3-mer. In this graph, the two distinct strings are not obvious because the *k*-mer is relatively small. There are many cycles within the graph, and the start and end *k*-mers for each string are obfuscated because of the structure. Algorithms that traverse this graph would need to explore every path in hopes of finding a bridge connecting two specific *k*-mers. This can lead to long computations or failure to find bridges if a branching limit is imposed on the traversal method. The corresponding 5-mer de Bruijn graph is show in (b). Every 5-mer from the two strings is a node in the graph. An edge indicates that the 4-mer suffix of the source 5-mer is the same as the 4-mer prefix of the destination 5-mer. In contrast to the 3-mer graph, the two distinct paths in this 5-mer graph are obvious because there are no shared 5-mers between the two strings. Algorithms that traverse this graph would find connecting *K*-mers to be a comparatively simple operation because there is no branching or cycles in the graph.

When the same sequences are viewed through a longer *K*-mer de Bruijn graph, the number of merged, ambiguous paths strictly decreases because an increasing amount of similarity is required for the paths to become merged in the graph. To demonstrate this idea, Figure 4b shows the 5-mer de Bruijn graph for the same two strings from Figure 4a. Note that while traversal of the graph for 3-mers was relatively difficult due to branching and cycles, the 5-mer graph is easily traversed because it is two disjoint linear paths. In practice, the short *k*-mer is still long enough to uniquely identify most areas of the genome. However, genomic characteristics such as low-complexity sequence, gene families, or repeat regions are difficult to traverse using short *k*-mers. Thus, our method uses the larger *K*-mer to bridge weak regions composed of repeated or low-complexity sequences that are too difficult to traverse using a small *k*-mer.

Because our method uses two passes with different sizes of *k* and *K*, it allows for less branching when *k* is small and more branching when *K* is large. As we just described, a small *k*-mer de Bruijn graph will have more branches and may require more computation to do a full depth-first traversal in repeat regions. To avoid this, we place more restrictions on the short *k*-mer traversals such that it primarily fixes the “easy” errors caused by sequencing. As a result, the “harder” traversals are pushed off to the long *K*-mer de Bruijn graph. The longer *K*-mer de Bruijn graph is less likely to have many branches and more likely to have a single pass connecting a start and target *K*-mer. In order to enable this approach, our method calculates a branch limit that scales linearly with the selected *k/K* value for the pass.

### 5.6 FMLRC Parameter Selection

FMLRC allows for four main parameters to be defined by the user: *T, F, k*, and *K. T* is the absolute minimum frequency required for a *k*-mer to be considered solid in the de Bruijn graph. *F* is the fraction of the median counts required for a *k*-mer to be considered solid in the de Bruijn graph. In all test cases, we used *T* = 5 and *F* =.10.

The second two parameters are the choice of *k* and *K* for the short and long correction passes. To gain some insight into what values of *k* and *K* are best, we ran multiple tests using the *E. coli* K12 MG1655 dataset. We allowed *k* = [17,19,21,23,25] and *K* = [–, 49, 59, 69, 79, 89], leading to a total of 30 test cases. The test cases with *K* = – indicate that no second *K*-mer pass was performed (it is only using a one-pass, short *k*-mer for correction). For each test case, we ran FMLRC, aligned the corrected reads to the reference genome, and then gathered statistics on the resulting alignment. We counted the the total number of bases that matched the reference genome and the total edit distance of the mappings. We also calculated the percentage of bases that matched the reference genome by taking the total number of matching bases and dividing by the total number of bases in the corrected reads. The results of this experiment are shown in Table 3.

**Table 3:**
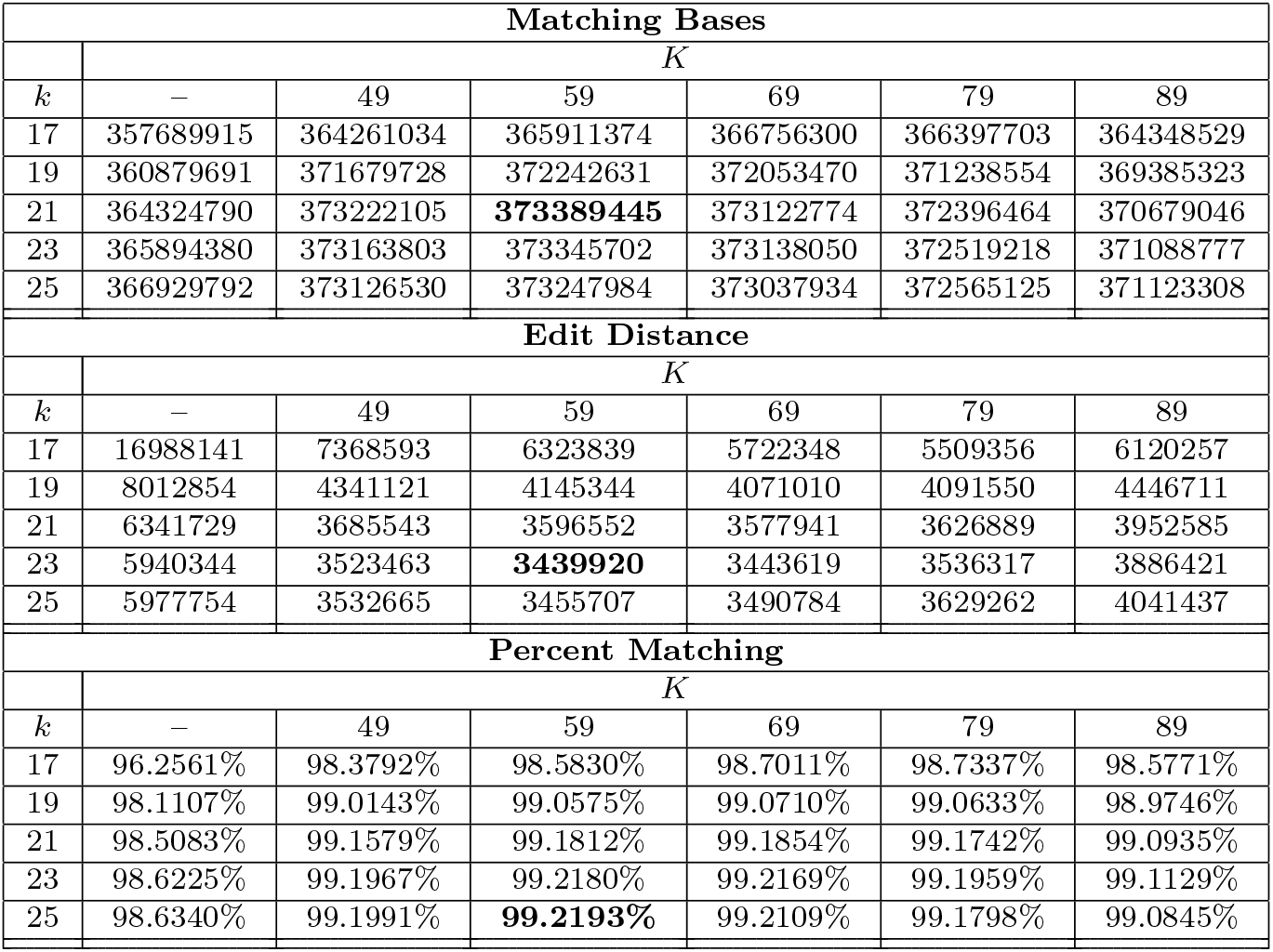
Choosing *k* and *K*. This table shows the result of running FMLRC using many different values for *k* and *K* for an *E. coli* dataset. The test cases with *K* = – indicate that no second pass of correction using the long *K*-mer was performed, so those test cases use a single pass short *k*-mer only. After correcting the reads, we aligned the results using BLASR and gathered statistics on the alignments. Matching bases indicates the number of matching bases across all mappings. Edit distance indicates the total number of mismatches, insertions, and deletions across all mappings. Percent matching is calculated as the total number of matching bases divided by the total number of bases that aligned. For each statistic, the best result is bolded in the above table. To summarize, a *K* = 59 is the column containing the test cases with the largest matching bases, lowest edit distance, and highest percent matching. Additionally, all tested values of *K* for a long *K*-mer pass improves the results over a single *k*-mer pass.

For each statistic measured, the best result used *K* = 59, but three different values for *k*. In all of our tests, performing a second pass with the long *K*-mer always improved all three statistics. Additionally, these results show that using a *K*-mer that is too large can have a negative impact on the performance of the correction. If the size of the *K*-mer relative to the read is too large, then the number of nodes in the de Bruijn graph will be reduced leading to difficulties during graph traversal. In our tests, while *K* = 59 is usually the best, *K* = 49 or *K* = 69 generally have similar performances. In contrast, the difference in edit distance from 69 to 79 and from 79 to 89 is more dramatic, indicating that there may be too few 79-mers and 89-mers with enough coverage to correct the reads. It is worth noting that the “best” *k* and *K* is likely data-dependent because differences in coverage, sequencing quality, and sequencing content will impact the ability of FMLRC to find solid *k*-mers and perform corrections.

### 5.7 Long read correction comparison

We compare FMLRC v0.1.2 to the LoRDEC v0.6 hybrid correction method [22]. This method corrects reads using a short *k*-mer, pruned de Bruijn graph. For all tests, we ran LoRDEC with the options `-k 21 -s 5` indicating a *k*-mer of length 21 and that any *k*-mer with frequency less than 5 is pruned from the de Bruijn graph. We also compared to a combined approach where LoRDEC is used for the short *k*-mer correction and FMLRC is used for the long *K*-mer correction.

For each dataset, we selected *k* and *K* based on the length and coverage of reads in the short-read sequencing datasets. For all test cases, we used *k* = 21 for both LoRDEC and FMLRC. For K, we used 59 for the three organisms with smaller genomes (< 25 million basepairs) and 69 for the *A. thaliana* dataset because it has a larger genome (> 100 million basepairs). We tested LoRDEC(k), LoRDEC(k) combined with FMLRC(K), and FMLRC(*k, K*) correction methods.

After running each correction method, we aligned the resulting FASTA file to the corresponding reference genome for the organism using BLASR v2.0.0 [8]. For BLASR, the only extra parameter we used was `-hitPolicy randombest` to force the alignment to keep only the best mapping for each read. Note that BLASR will also split reads into multiple subreads if it detects multiple subreads that best align to different genomic locations.

### 5.8 Performance Testing

Performance tests were run on a single machine running Ubuntu 14.04 with 32 GB memory and an Intel Xeon E5-2620 6-core 2.00 GHz processor. The machine is connected to a 1 TB HDD for reading and writing any necessary input or output files. For measuring performance, we used the built-in `/usr/bin/time` function to extract real time, user time, memory usage, and CPU utilization. Each method was allowed eight processes for computing the results.

### 5.9 Data Description

We tested the correction algorithms on five publicly available PacBio datasets. The PacBio datasets were downloaded for *E. coli* K12 MG1655, *P. falciparum* 3d7, *S. cerevisiae* W303, and *A. thaliana* Ler-0[5]. For each dataset, we also downloaded a publicly available short-read sequencing dataset: a publicly available dataset[24] for *E. coli*, SRR1503358 from Genbank for *P. falciparum*, SRR1652473 for *A. thaliana*, and publicly available data[23] for *S. cerevisiae*.

### 5.10 *De novo* assembly comparison

We ran the long-read correction and *de novo* assembly methods listed in Table 1 on the *E. coli*, *S. cerevisiae*, and *A. thaliana* datasets listed above. All were run on Killdevil, a heterogeneous Linux-based cluster with more than 9600 cores and 48Gb - 1Tb RAM per node at Research Computing at UNC. All jobs had a hard limit of 16 processes and one week wall-clock run time. For larger genomes such as *A. thaliana*, several methods including hybridSPAdes, canu, and ECTools failed after exceeding these limits or exceeding 1Tb main memory.

We used Quast v4.1 [13] to assess these assemblies using default parameters.

## 6 Data Access

FMLRC is a publicly available C++ program^3^. The implementation requires that a BWT of the short-read dataset has already been constructed and is in the run-length encoded format of the msbwt package^4^.

## 7 Acknowledgements

This work was supported in part by funding from the National Science Foundation (C.D.J. DEB-1457707), North Carolina Biotechnology Center (C.D.J, 2013-MRG-1110), and the University Cancer Research Fund (C.D.J.).

## Footnotes

1 https://github.com/holtjma/msbwt/wiki/Converting-to-msbwt’s-RLE-format

2 https://github.com/holtjma/msbwt/wiki/Converting-to-msbwt’s-RLE-format

3 http://github.com/holtjma/fmlrc

4 http://github.com/holtjma/msbwt

## References

[1] Dmitry Antipov, Anton Korobeynikov, Jeffrey S. McLean, and Pavel A. Pevzner. hybridspades: an algorithm for hybrid assembly of short and long reads. Bioinformatics, 32(7):1009–1015, 2016.

[2] Kin Fai Au, Jason G Underwood, Lawrence Lee, and Wing Hung Wong. Improving pacbio long read accuracy by short read alignment. PLoS One, 7(10):e46679, 2012.

[3] Markus J Bauer, Anthony J Cox, and Giovanna Rosone. Lightweight bwt construction for very large string collections. In Combinatorial Pattern Matching, pages 219–231. Springer, 2011.

[4] Konstantin Berlin, Sergey Koren, Chen-Shan Chin, James P Drake, Jane M Landolin, and Adam M Phillippy. Assembling large genomes with single-molecule sequencing and locality-sensitive hashing. Nat Biotech, 33(6):623-630, 06 2015.

[5] Pacific Biosciences. Pacbio datasets. https://github.com/pacificbiosciences/devnet/wiki/datasets.

[6] de NG Bruijn. A combinatorial problem. Proceedings of the Koninklijke Nederlandse Akademie van Wetenschappen. Series A, 49(7):758, 1946.

[7] Michael Burrows and David J Wheeler. A block-sorting lossless data compression algorithm. 1994.

[8] Mark J Chaisson and Glenn Tesler. Mapping single molecule sequencing reads using basic local alignment with successive refinement (blasr): application and theory. BMC bioinformatics, 13(1):238, 2012.

[9] Shigang Wu Jue Ruan Zhanshan Ma Chengxi Ye, Chris Hill. Dbg2olc: Efficient assembly of large genomes using long erroneous reads of the third generation sequencing technologies. arXiv:1410.2801, May 2016.

[10] Chen-Shan Chin, David H Alexander, Patrick Marks, Aaron A Klammer, James Drake, Cheryl Heiner, Alicia Clum, Alex Copeland, John Huddleston, Evan E Eichler, et al. Nonhybrid, finished microbial genome assemblies from long-read smrt sequencing data. Nature methods, 10(6):563–569, 2013.

[11] Paolo Ferragina and Giovanni Manzini. An experimental study of an opportunistic index. In Proceedings of the twelfth annual ACM-SIAM symposium on Discrete algorithms, pages 269–278. Society for Industrial and Applied Mathematics, 2001.

[12] Seth Greenstein, James Holt, and Leonard McMillan. Short read error correction using an fm-index. In Bioinformatics and Biomedicine (BIBM), 2015 IEEE International Conference on, pages 101–104. IEEE, 2015.

[13] Alexey Gurevich, Vladislav Saveliev, Nikolay Vyahhi, and Glenn Tesler. Quast: quality assessment tool for genome assemblies. Bioinformatics, 29(8):1072–1075, 2013.

[14] Shinjae Yoo Shoshana Marcus W. Richard McCombie Michael Schatz Hayan Lee, James Gurtowski. Error correction and assembly complexity of single molecule sequencing reads. June 2014.

[15] James Holt and Leonard McMillan. Merging of multi-string bwts with applications. Bioinformatics, page btu584, 2014.

[16] Sergey Koren, Michael C Schatz, Brian P Walenz, Jeffrey Martin, Jason T Howard, Ganeshkumar Ganapathy, Zhong Wang, David A Rasko, W Richard McCombie, Erich D Jarvis, et al. Hybrid error correction and de novo assembly of single-molecule sequencing reads. Nature biotechnology, 30(7):693–700, 2012.

[17] Heng Li. Fast construction of fm-index for long sequence reads. Bioinformatics, page btu541, 2014.

[18] Heng Li. Minimap and miniasm: fast mapping and de novo assembly for noisy long sequences. Bioinformatics, 32(14):2103–2110, 2016.

[19] Nicholas J Loman, Joshua Quick, and Jared T Simpson. A complete bacterial genome assembled de novo using only nanopore sequencing data. Nat Meth, 12(8):733-735, 08 2015.

[20] Mari Miyamoto, Daisuke Motooka, Kazuyoshi Gotoh, Takamasa Imai, Kazutoshi Yoshitake, Naohisa Goto, Tetsuya Iida, Teruo Yasunaga, Toshihiro Horii, Kazuharu Arakawa, Masahiro Kasahara, and Shota Nakamura. Performance comparison of second- and third-generation sequencers using a bacterial genome with two chromosomes. BMC Genomics, 15(1):1–9, 2014.

[21] Gene Myers. Daligner. https://github.com/thegenemyers/daligner.

[22] Leena Salmela and Eric Rivals. Lordec: accurate and efficient long read error correction. Bioinformatics, page btu538, 2014.

[23] Michael Schatz. Schatz lab data. http://schatzlab.cshl.edu/data/ectools/.

[24] SPAdes. Standard isolate e. coli. http://spades.bioinf.spbau.ru/.

[25] Todd J Treangen and Steven L Salzberg. Repetitive dna and next-generation sequencing: computational challenges and solutions. Nature Reviews. Genetics, 13(1):36-46, 11 2011.

[26] Son Pham Vineet Bafna Viraj Deshpande, Eric DK Fung. Cerulean: A hybrid assembly using high throughput short and long reads. arXiv:1307.7933, July 2013.

